# The chicken or the egg? Plastome evolution and a novel loss of the inverted repeat in papilionoid legumes

**DOI:** 10.1101/2021.02.04.429812

**Authors:** Chaehee Lee, In-Su Choi, Domingos Cardoso, Haroldo C. de Lima, Luciano P. de Queiroz, Martin F. Wojciechowski, Robert K. Jansen, Tracey A Ruhlman

## Abstract

The plastid genome (plastome), while surprisingly constant in gene order and content across most photosynthetic angiosperms, exhibits variability in several unrelated lineages. During the diversification history of the legume family Fabaceae, plastomes have undergone many rearrangements, including inversions, expansion, contraction and loss of the typical inverted repeat (IR), gene loss and repeat accumulation in both shared and independent events. While legume plastomes have been the subject of study for some time, most work has focused on agricultural species in the IR-lacking clade (IRLC) and the plant model *Medicago truncatula*. The subfamily Papilionoideae, which contains virtually all of the agricultural legume species, also comprises most of the plastome variation detected thus far in the family. In this study 33 newly sequenced plastomes of papilionoid legumes and outgroups were evaluated, along with 34 publicly available sequences, to assess plastome structural evolution in the subfamily. In an effort to examine plastome variation across the subfamily, just ∼20% of the sampling represents the IRLC with the remainder selected to represent the early-branching papilionoid clades. A number of IR-related and repeat-mediated changes were identified and examined in a phylogenetic context. Recombination between direct repeats associated with *ycf2* resulted in intraindividual plastome heteroplasmy. Although loss of the inverted repeat has not been reported in legumes outside of the IRLC, one genistoid taxon was found to completely lack the typical plastome IR. The role of the IR and non-IR repeats in driving plastome change is discussed.

**Significance statement:** Comparative genomic approaches employing plastid genomes (plastomes) have revealed that they are more variable across angiosperms than previously suggested. This study examined 64 species of Fabaceae and outgroups, including 33 newly sequenced taxa, to explore plastome structural evolution of the subfamily Papilionoideae in a phylogenetic context. Several unusual features of the inverted repeat highlight the importance of recombination in plastomic structural changes within and between individuals and species.

## Introduction

The plant cell is endowed with three genetic compartments, the nucleus, the mitochondrion and the plastid. Plastids uniquely possess pluripotency, that is, they are capable of interconversion between types depending on developmental and environmental cues. The most widely recognized of these types are the chlorophyll-containing chloroplasts of photosynthetic cells. The plastid genome (plastome) includes the entire DNA content in all plastid types of an individual and comprises many copies of the unit genome or plastome monomer, defined as one full gene complement, along with intergenic regions. Typically each monomer contains two single copy (SC) regions defined by their length, large and small (LSC and SSC, respectively), separated by a large inverted repeat (IR; typically ∼25 kb). The plastomes of photosynthetic angiosperms commonly encode around 80 proteins, 30 tRNAs and four rRNAs, all of which function in the organelle in conjunction with nuclear-encoded proteins imported from the cytosol (Jansen and Ruhlman, 2012; Ruhlman and Jansen, 2021).

Although the typical plastome monomer is only around 150 kb in length, the plastome may represent up to 20% of all the DNA found in a photosynthetically active cell due to its highly iterative nature (Bendich, 1987; Rauwolf *et al*., 2010). Early clues suggested an assemblage of circular molecules each containing one to several copies of the unit genome, a view likely influenced by the endosymbiotic origins of the organelle (Sagan, 1967; Kolodner and Tewari 1972; Herrmann *et al*., 1975; Bedbrook and Kolodner, 1979). The current understanding points to predominantly linear and branched molecules comprising covalently linked repeating units, segments of which undergo recombination both within and between units and molecules (Bendich and Smith, 1990; Lilly *et al*., 2001; Oldenburg and Bendich, 2004; Oldenburg and Bendich, 2015; Oldenburg and Bendich, 2016). Both the LSC and SSC undergo inversion as a result of recombination between IR copies in different monomeric units. The resulting population of monomers likely exists as at least four roughly equivalent isomeric units that vary in their junctions between single copy and IR seqeunces. Both early (Palmer, 1983; Palmer *et al*., 1987) and more recent (Oldenburg and Bendich, 2004; Blazier *et al*., 2016a) investigators have noted the similarity of this phenomenon to the equimolar isomers of the HSV-1 genome, which also contains SC and IR regions.

The typical structure of contemporary plastome units, two single copy regions separated by an inverted repeat, is thought to be ancestral based on its widespread presence across the phylogeny of photosynthetic eukaryotes (Ruhlman and Jansen, 2021; Mower and Vickrey, 2018). Despite years of speculation as to the origin and purpose of this arrangement, particularly with respect to the function of the plastome IR, a satisfying explanation remains elusive. Among the earliest hypotheses was the notion that the IR played a role in maintaining plastome structural integrity. In land plants, IR presence seemed to correlate with the conservation of gene order (Palmer and Thompson, 1982), however as more plastomes with and without the typical IR have been examined this suggestion has become less appealing. Furthermore, while it is likely that the large plastome repeats are involved in replication initiation, the IR is not necessary to this essential function (Mühlbauer *et al*., 2002; Scharff and Koop, 2007).

Typical IRs also maintain the homogeneity of the sequences encoded in each copy. Genes situated in the IR tend to accumulate substitutions more slowly than those of the single copy regions (Birky and Walsh, 1992; Maier *et al*., 1995; Perry and Wolfe, 2002; Guisinger *et al*., 2010) and both copies are thought to be identical in nucleotide sequence. This suggests that gene conversion works to homogenize the repeat and this mechanism has been invoked to explain copy correction in plastid transformants when point mutations or foreign sequence are introduced into the IR (Ruhlman and Jansen, 2021).

Across angiosperms, IR extent ranges dramatically from complete absence to more than 80 kb (Chumley *et al*., 2006; Weng *et al*., 2014; Weng *et al*., 2017; Sanderson *et al*., 2015; Blazier *et al*., 2016a; Ruhlman *et al*., 2017; Mower and Vickrey, 2018; Solórzano *et al*., 2019; Lee *et al*., 2020; Li *et al*., 2020) and can be a major contributor to overall plastome expansion. Although IR expansion is more commonly associated with IR/SC junction migration, which includes or excludes sequence (Mower and Vickrey, 2018; Ruhlman and Jansen, 2021), occasionally insertion of foreign sequence (Goremykin *et al*., 2009; Iorizzo *et al*., 2012; Ma *et al*., 2015; Rabah *et al*., 2017; Burke *et al*., 2018) and accumulation of dispersed repeats and sequence deletions have been reported within the IR. For example, the largest gene within typical IRs is *ycf2*, encoding an ATPase motor protein involved in trans-membrane import of proteins translated in the cytoplasm (Shinozaki *et al*., 1986; Drescher *et al*., 2000; Kikuchi *et al*., 2018). The full length gene in typical IRs exceeds 8 kb, however the open reading (ORF) frame varies widely and has been independently lost from the plastomes of several lineages (Guisinger *et al*., 2010; Ruhlman and Jansen, 2018; Shrestha *et al*., 2019; Jin *et al*., 2020a,b; Lee *et al*., 2020), contributing to IR length variation across photosynthetic angiosperms.

Variation in IR extent, IR involvement in replication initiation and efficient gene conversion are all intimately connected to recombination. Recombination within and between plastome monomers has been a subject of speculation and examination nearly as long as the plastome has been studied. By the 1970s, groups using various approaches identified the IR in several taxa and postulated that recombination between the repeated sequences reversed the orientation of the region that separated the two copies. It was also noted that not all plastomes, pea (*Pisum sativum*) for example, contained a large IR, which represented up to ∼30% of the unit genome when present among the examined taxa (Bedbrook and Bogorad, 1976; Bedbrook and Kolodner, 1979) and that recombination between the repeats would tend to prevent divergence of IR sequences (i.e. gene conversion; Kolodner and Tewari, 1979).

Indeed, recombination both within and between unit genomes can not only homogenize but also diversify the plastome, whether mediated by IR sequences, other repeats or single copy sequences. Typical plastomes, those that exhibit no irregularities in substitution rates or structural arrangements, likely exist as a population of molecules comprising predominantly linear and branched concatenations of monomers as well smaller linear and circular forms. Isomeric units each contain the full complement of plastome sequences but differ with respect to the relative orientation of their single copy regions, much as predicted by early observations. Recombination between non-homologous repeats (i.e. ‘copy A’ in one unit genome aligns with ‘copy B’ in another unit) is analogous to IR mediated recombination that reverses the polarity of single copy regions and can lead to gross rearrangements such as inversions, deletions and duplications. In some cases, although not stoichiometrically maintained, subgenomes with sequence arrangements that differ from the predominant plastome monomer persist (Guo *et al*., 2014; Ruhlman *et al*., 2017; Lee *et al*., 2020).

Inversion of large plastome blocks have provided useful phylogenetic markers (Jansen and Palmer, 1987; Bruneau *et al*., 1990; Lavin *et al*., 1990; Downie and Palmer, 1992; Raubeson and Jansen, 1992; Doyle *et al*., 1996; Jansen *et al*., 2008; Martin *et al*., 2014; Rabah *et al*., 2018) as these rearrangements tend to exhibit lower levels of homoplasy than gene and intron losses (Raubeson and Jansen, 2005). Many have noted the presence of repeats flanking inverted blocks and suggested that these repeats played a role in reversing the polarity of the affected segment (Howe *et al*., 1988; Lee *et al*., 2007; Guisinger *et al*., 2011). Ancient examples, such as in vascular plants (Raubeson and Jansen, 1992) and Asteraceae (Jansen and Palmer, 1987) demonstrate that inversions present a robust phylogenetic signal. However, repeat mediated inversion will not always be stable, and inverted segments may revert their orientation with time, likely by the same dynamic mechanism that generates isomeric units of the plastome. The 36 kb inversion first described by Martin et al. (2014) could be such a case. The inversion in the genistoid legume *Lupinus* was flanked by *trnS* sequences and was described as specific to the core genistoid (Martin *et al*., 2014). Subsequent studies have revealed the presence of the 29 bp *trnS*-associated inverted repeat across Fabaceae and identified another incidence of inversion in the robinioid taxon *Robinia pseudoacacia* (Schwarz *et al*., 2015).

With 765 genera and almost 20,000 species, the legume family (Fabaceae or Leguminosae) represents ca. 7% of flowering plant diversity. Currently the family includes six subfamilies (LPWG 2017) that evolved almost simultaneously at the Cretaceous-Paleogene boundary (Koenen et al. 2020a, 2020b). Papilionoid legumes (Fabaceae, subfamily Papilionoideae) comprise an estimated 503 genera and 14, 000 species and are extraordinarily diverse in ecology and biogeographical range (Schrire *et al*., 2005; LPWG 2017). The subfamily contains all the culinary pulses, beans and peas, in addition to the important forage and feed crop, alfalfa (*Medicago sativa*). With few exceptions, the most familiar of the legume crops belong to the IR-lacking clade (IRLC; Lavin *et al*., 1990; Wojciechowski *et al*., 2004). Comparison between pea and mung bean (*Vigna radiata*; Palmer and Thompson, 1981), revealed that homologous sequences in the IR-lacking pea plastome were rearranged relative to mung bean, which includes an ∼23 kb IR. The authors noted the finding that pea plastome lacked the large repeat could suggest that the IR was an evolutionary relic, i.e having no contemporary function. A causal relationship between IR loss and dysregulation of structural homeostasis was later proposed following examination of pea and broad bean (*Vicia faba*; Palmer *et al*., 1983). Further comparisons revealed that the loss of the IR had not led to the rearrangement of the alfalfa and *Wisteria* plastomes (Palmer et al., 1987). Numerous plastomes have now been examined that contain IRs ranging widely in size and other examples that lack the large IR entirely, however no robust correlation to genome stability can be ascribed to its presence, absence or extent (Guisinger *et al*., 2011; Weng *et al*., 2014; Blazier *et al*., 2016a; Ruhlman and Jansen, 2018; Ruhlman and Jansen, 2021)

Plastome IR loss tends to carry a strong phylogenetic signal, that is, the losses are typically restricted to discrete, independent clades, having occurred once in a common ancestor of each lineage. Previous investigations of IRLC legumes have included closely related IR containing taxa for comparison and often included few representatives from either group. In this study 33 newly sequenced plastomes of papilionoid legumes and outgroups were evaluated, along with GenBank data, to assess plastome structural evolution in the subfamily. Approximately 20% of included plastomes were from IRLC species, while the remaining proportion are distributed across the subfamily and outgroups. This study examined 64 species of Fabaceae and outgroups, including 33 newly sequenced taxa, to explore plastome structural evolution of the subfamily Papilionoideae in a phylogenetic context. Several unusual features of the inverted repeat highlight the importance of recombination in plastomic structural changes within and between individuals and species.

## Materials and Methods

### Taxon sampling and DNA extraction

A total of 67 taxa were selected, including 59 Papilionoideae from previously recognized clades (Cardoso *et al*., 2012), five taxa from other subfamilies of Fabaceae along with three additional Fabales outgroup taxa from Polygalaceae, Surianaceae and Quillajaceae. Plastomes for 34 taxa were downloaded from NCBI (https://www.ncbi.nlm.nih.gov/genome/organelle/) (Table S1).

For the 33 newly sequenced taxa, plants were either grown from seed obtained from the Desert Legume Program (University of Arizona, Tuscon) in the UT-Austin greenhouse or sampled from field sites in Brazil and Texas (Table S1). Emergent leaves were collected from young plants or from the actively growing tips of field specimens. Collected leaves were flash frozen in liquid nitrogen and stored (as needed) at −80°C for DNA isolation. Voucher specimens of the greenhouse-grown taxa and Texas field collections were deposited in the Billie L. Turner Plant Resource Center (TEX-LL) at UT-Austin. Voucher specimens of 29 taxa were deposited in the Herbário da Universidade Estadual de Feira de Santana (HUEFS), Bahia, Brazil, or may be found in the Rio de Janeiro Botanical Garden Herbarium (RB). All vouchers include the sample ID from Table S1. Total genomic DNA was extracted using the NucleoSpin Plant II, Mini Kit for DNA from plants (Macherey-Nagel, Düren, Germany) or by the method of Doyle and Doyle (1987) using 3% polyvinylpyrrolidone in the extraction buffer (see †, Table S1). All isolated samples were stored at −20 °C.

### DNA sequencing, assembly and annotation

Total genomic DNA for 33 taxa was shipped to the Beijing Genomics Institute (BGI; Shenzhen) and sample quality was confirmed. Libraries with insert sizes of ∼350 bp were prepared and sequenced generating a minimum of 20 million, 150 bp paired-end (PE; minimum 40 million reads total) reads using the BGI DNBseq^™^ platform.

All quality filtered reads (Table S1) were assembled *de novo* using Velvet version 1.2.08 (Zerbino and Birney, 2008) with multiple k-mers at the Texas Advanced Computing Center following Lee et al. (2020). Assembled plastid contigs were imported into Geneious Prime (https://www.geneious.com) and all protein-coding, tRNA and rRNA genes were annotated using BLAST against the reference plastome of *Cercis glabra* (NC_036762). Annotated contigs were assembled in Geneious Prime by trimming overlapping regions of each contig to produce draft unit genomes. The untrimmed contigs used to assemble the draft were then aligned with the draft unit genomes to validate IR boundaries and the whole plastome sequences. The PE reads were mapped to the completed plastome sequences using Bowtie 2 (Langmead *et al*., 2009) to assess the depth of coverage for whole plastome sequences.

Annotations of all protein-coding, tRNA and rRNA genes of *Parkinsonia aculeata* were manually examined and corrected in Geneious by comparing with published reference plastomes of Fabaceae downloaded from NCBI (https://www.ncbi.nlm.nih.gov/genome/organelle/) and used as a reference to confirm annotations for all the genes in the complete plastomes of remaining 32 taxa that were newly generated in this study.

### Plastome structural analysis

Whole-genome alignments were carried out using Mauve v.2.3.1 in Geneious Prime with the progressiveMauve algorithm and default parameters (Darling *et al*., 2010). The complete plastomes of 64 legume taxa were each aligned with a reference plastome, *Suriana maritima*, which has the angiosperm ancestral plastome gene order (Ruhlman and Jansen, 2021). Each plastome showing changes in gene order relative to the reference was individually aligned with the reference. Locally colinear blocks (LCBs) were manually numbered and breakpoint (BP) and reversal distances were calculated using the web-based application CREx (Common Interval Rearrangement Explorer; Bernt *et al*., 2007) with default parameters. Complete plastomes were edited to exclude one repeat copy from IR-containing plastomes prior to Mauve alignment. The inversion of the 50 kb-inversion clade (Doyle *et al*., 1996; Cardoso *et al*., 2012) was not considered in this assessment. The *Wisteria* plastome, which is a relatively unrearranged (single inversion relative to reference *Suriana*) and an early diverging IRLC taxon, was aligned with *C. scadens* using progressiveMauve to assess relative plastome organization between two IR-less plastomes.

Plastome assemblies for four taxa, *Harleyodendron unifoliolatum, Diplotropis ferruginea, Poiretia bahiana* and *Galactia jussiaeana*, were edited to create maps of hypothetical plastome haplotypes that could arise from recombination between direct repeats situated near the start of the Ycf2 coding region. Quality-filtered paired-end reads (insert size up to 700 bp) were mapped to the hypothetical haplotype drafts using Bowtie 2. The PE reads that were congruent only with hypothetical haplotypes were counted.

To confirm the presence of alternate plastome *ycf2* haplotypes, PCR was performed using total genomic DNA of four taxa with nine oligonucleotide primers (Table S2) designed using Primer 3 (Untergasser *et al*., 2012; Table S2) in Geneious Prime. PCR reactions were carried out in 13 ul volumes and included 4.5 ul distilled water, 6.5 ul FailSafe^™^ PCR 2X PreMix J (Lucigen, Middleton), 0.5 ul Taq DNA polymerase, 0.5 ul of each primer (10 pmole/ul), and 0.5 ul genomic DNA (∼40 ng). Different combinations of primers were used to amplify variable target regions. Reactions included an initial denaturation step at 98 °C for 1 min, 35 cycles of denaturation at 98 °C for 30 sec, annealing at 55 °C for 30 sec and extension at 72 °C for 1 min, and final extension at 72 °C for 5 min. Amplification products were evaluated in 2 % agarose with Low Molecular Weight DNA Ladder (New England BioLabs, Ipswich). Residual primers and nucleic acids were removed from PCR reactions with Exonuclease I (New England BioLabs) and FastAP Thermosensitive Alkaline Phosphatase (Thermo Scientific, USA) and PCR products were Sanger sequenced at the Genome Sequencing and Analysis Facility at UT-Austin.

### Repetitive DNA sequence analysis

Repeat content for all 67 taxa was estimated by enumerating tandem and dispersed repeats. For repeat analyses, one copy of the IR was removed from IR-containing taxa. Tandem repeats were identified using Tandem Repeats Finder v.4.09 with default options (Benson, 1999). Dispersed repeats were analyzed by the command line version BLAST v.2.8.1+ (Altschul *et al*., 1990) using each plastome as both query and subject with a word size of 16 and percent identity of 80%. All BLAST hits were retained. Sequence coordinates of both tandem and dispersed repeats were transferred to each plastome in Geneious and the percentage of repetitive DNA was calculated. The Pearson correlation test was carried out to evaluate the relationship between the proportion of dispersed repeats and reversal distance for each taxon.

### Phylogenetic analysis

All 73 protein-coding genes (Table S3) shared among plastomes of all 64 legume taxa and the outgroups *Polygala fallax* (MT762166), *Suriana maritima* (NC_047313) and *Quillaja saponaria* (NC_047356) were extracted in Geneious Prime. All protein-coding sequences were aligned individually for each gene using MAFFT (Katoh and Standley, 2013) and concatenated into a single aligned data set. Before and after MAFFT alignments, all gene sequences and aligned data sets were manually inspected to verify the reading frame and refine poorly aligned regions. The evolutionary substitution models and best partition scheme were determined using software IQ-TREE v.1.6.12 (Nguyen *et al*., 2015; Chernomor *et al*., 2016) to identify the best-fit models of substitution for each partition. IQ-TREE was implemented to perform maximum likelihood (ML) analysis to reconstruct the phylogenetic tree and assess branch support with the ultra-fast bootstrap option with 2000 pseudoreplicates. The ML tree with bootstrap support values was visualized using FigTree v.1.4.3 (http://tree.bio.ed.ac.uk/software/figtree/).

## Results

### An independent IR loss in *Camoensia scadens*

Sequencing, assembly and read mapping experiments confirmed that the complete plastome of the genistoid legume *Camoensia scandens* lacked the canonical IR, resulting in the plastome size of 127,752 bp (Table S1), whereas plastomes of other taxa in closely related clades all included the typical IR (Figure 1; Figure S1; Table S1). Two overlapping sequences of 107 bp and 35 bp were detected in the region downstream of *ndhF* in reverse orientation in the upstream and downstream regions of *rpl16*. (Figure 2A). Quality-filtered reads mapped to the complete plastome of *Camoensia* confirmed that they were evenly distributed across the entire plastome assembly supporting the loss of one IR coy (Figure 2B). Whole genome alignment to the early branching IRLC legume *Wisteria* did not support gross reorganization of the *Camoensia* plastome (Figure S2). Three inversions were noted, however the timing of these changes, prior to or post IR deletion, is unknown.

**Figure 1:**
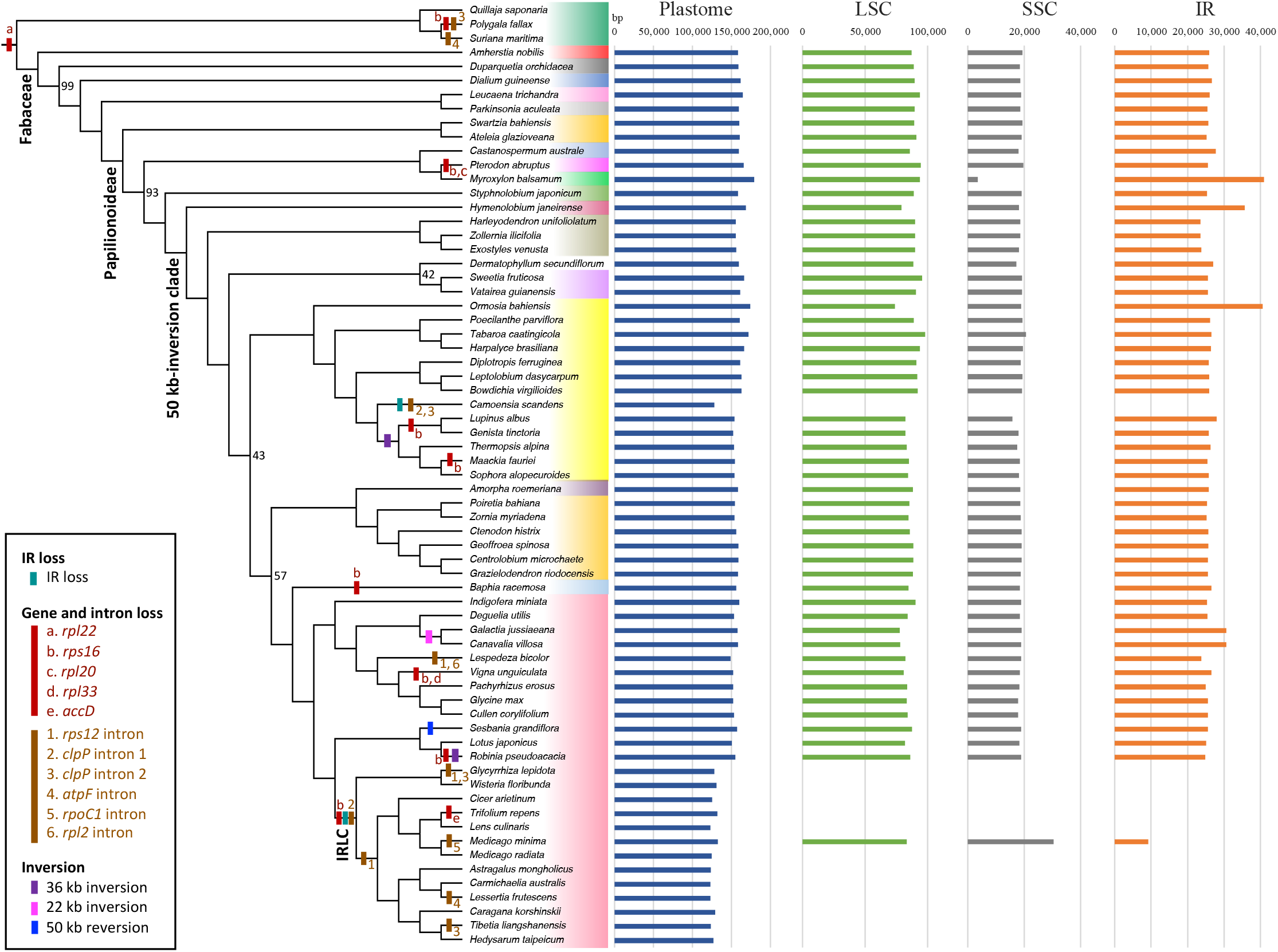
Phylogenetic relationships and distribution of plastome variation in Papilionoideae. The cladogram is derived from the phylogram shown in Figure S1. The legend (lower left) indicates selected plastome changes, which are marked on the relevant branches. Taxon names are contained in shaded blocks according to clades, color scheme is repeated in Figure S1 to indicate clade names. Bootstrap values less than 100% are shown at the nodes, except for five terminal nodes, which are shown on the phylogram in Figure S1. Plastome size variation is indicated in basepairs (bp; see also Table S1). LSC, large single copy region; SSC, small single copy region; IR, inverted repeat.

**Figure 2:**
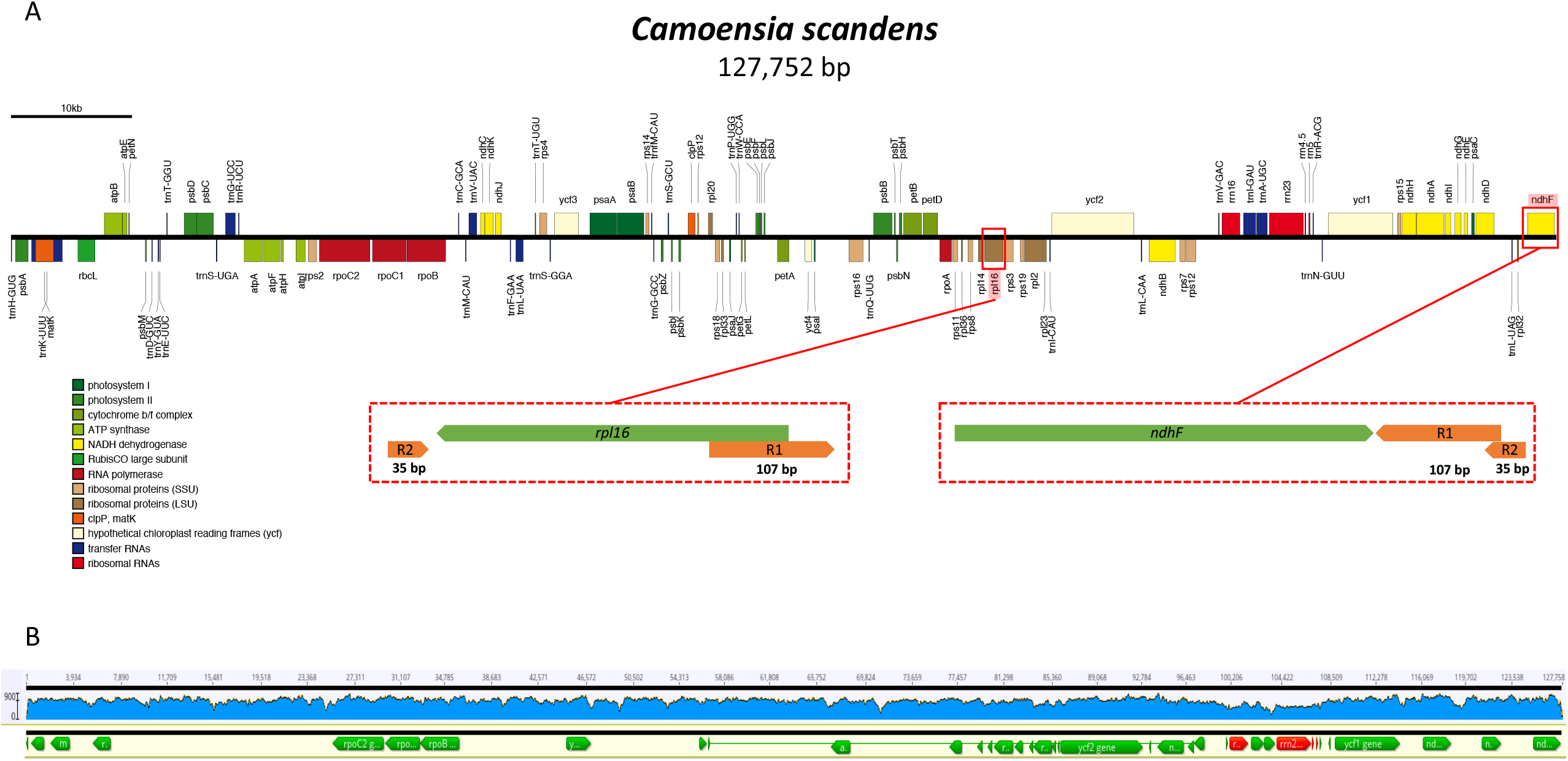
The *Camoensia scadens* plastome lacks the large inverted repeat. The schematic representation of the *C. scadens* plastome monomer (A) is drawn to scale (bar at upper left) and includes gene blocks colored to indicate functional groups (see legend, left). Repeated sequences (R1, R2) flanking the remaining, single copy inverted repeat (IR) sequences are highlighted. Sequencing reads mapped to the monomer assembly (B) are indicated by blue histograms and depth of coverage is given by the scale at left. Genome coordinates (above) are indicated above and selected gene blocks (below; green) are shown, including the *rrn* genes (red) typically found in the plastome IR. Basepairs, bp.

### Plastome organization, repeat accumulation and size variation across Papilionoideae

#### Gene and intron loss

Among all 67 taxa, there were 73 shared, unique protein-coding genes, with 76 or 77 annotated in each plastome. Several gene and intron losses were noted. All legume taxa shared the loss of *rpl22* and two additional independent losses of *rps16* gene were identified in *Baphia* and *Pterodon*. Other genes, *rpl20, rpl33* and *accD*, were psuedogenized or missing in the plastomes of *Pterodon, Vigna* and *Trifolium*, respectively (Figure 1; Table S1).

Papilionoideae plastomes showed three independent losses of the *rps12* intron. *Tibetia liangshanensis, Camoensia scadens* and *Glycyrrhiza lepidota* entirely lack *clpP* introns. Intron losses were also noted in *Medicago minima* and *Lessertia frutescens*, from *rpoC1* and *atpF*, respectively (Figure 1; Table S1).

#### Inversion

While none were noted among representatives of other Fabaceae subfamilies, a number of papilionoid species contained plastome inversions (Table S1). The shared 50 kb inversion (Doyle *et al*., 1996) was not counted as an event in the analysis, however a number of taxa have experienced reversion of large portions of this feature. The 36 kb inversion described by Martin et al. (2014) was detected in the *Lupinus/Genista* clade and was also present in the sister clade that included *Thermopsis, Maackia* and *Sophora*. Inversions of ∼ 22 kb shared by *Galactia* and *Canavalia*, and of ∼50 kb unique to *Hymenolobium*, each reverted a portion of the 50 kb inversion. A single taxon, *Sesbania grandiflora*, did not contain the 50 kb inversion and was essentially colinear with plastomes of early-branching papilionoid lineages, caesalpinioids and mimosoids, as well as the outgroup *Suriana* (Figure 1; Table S1).

#### Inverted repeat expansion and plastome size variation

The size of IR-containing plastomes varied from ∼132 kb to ∼179 kb with the LSC, SSC, IR ranging in size from approximately 74 kb to 98 kb, 3.5 kb to 30 kb, and 9 kb to 41 kb, respectively (Figure 1, Table S1). Expansion of the IR in four taxa, *Ormosia bahiensis, Hymenolobium janeirense, Galactia jussiaeana* and *Canavalia villosa* resulted in reduction of the LSC and overall plastome expansion. Reduction of the SSC as a result of IR expansion left just 3,584 bp of single copy sequence and two genes (*ndhF* and *rpl32*) in *Myroxylon balsamum* SSC. This expansion combined with an accumulation of tandem repeats in the LSC yielded the largest plastome among investigated legumes (>179 kb; Table S1). Repeat accumulation substantially expanded the LSC (98,188 bp) of *Tabaroa caatingicola*, where tandem repeats represented 10.34% of the plastome (171,932 bp).

#### Dispersed and tandem repeat accumulation

The proportion of dispersed and tandem repeats in legume plastomes ranged from 0.69 to 10.08 % and 0.69 to 10.34 %, respectively (Table S1). The greatest proportion of tandem repeats was observed in *Tabaroa caatingicola* followed by *Hedysarum taipeicum* (8.54 %), *Pterodon abruptus* (8.35 %), *Sweetia fruticosa* (7.5%) and *Myroxylon balsamum* (6.2 %). Whereas most legume plastomes contained few dispersed repeats, two IRLC taxa, *Hedysarum taipeicum* and *Trifolium repens*, accumulated substantially more dispersed repeats: 10.08 % and 9.03 %, respectively. There was no significant positive correlation between dispersed repeat accumulation and reversal distance (inversion; *R*=0.47, *p*<0.001).

#### Conserved repeats and ycf2 length variation

A direct repeat was detected in plastomes of the 50-kb inversion clade taxa (Figure S1, Table S1). In *Harleyodendron unifoliataum* one copy of the repeat was situated in the intergenic region between *rpl23* and *trnI-CAU* upstream of *ycf2* (nucleotides 91,042 – 91,325; R1) and the other within the *ycf2* ORF (92,991 – 93,277; R2)(Figure 3, S1, Table S1). Both copies of the repeat were included in the IR of plastomes that contain this structure. The repeats were situated at nearly identical positions across all taxa and varied in size in a clade-dependent manner (Figure S1, Table S1). Among taxa that have not undergone IR loss, the repeats ranged from 76 bp to 308 bp, with identities from 98.7 % to 100 %. Blast results indicated that a deletion had occurred in R1 of several taxa, while several others experienced tandem repeat accumulation within R1. With some exceptions, IRLC taxa and *Camoensia* plastomes contained repeats that were either smaller (33 bp to 90 bp) and/or broken into noncontiguous segments (Figure S1, Table S1). The two earliest diverging IRLC taxa, *Glycyrrhiza lepidota* and *Wisteria floribunda*, contained repeats similar in size to non-IRLC plastomes, 268 bp and 302 bp, respectively. While R1 was truncated in several IRLC taxa, in *Cicer arietinum* and *Medicago radiata* R2 length was more conserved at 117 bp and 198 bp, respectively. In *Hedysarum taipeicum* (IRLC), both R1 and R2 were situated in the intergenic region as ORF prediction indicated an internal stop codon in *ycf2*, truncating the 5’ end of the gene.

**Figure 3:**
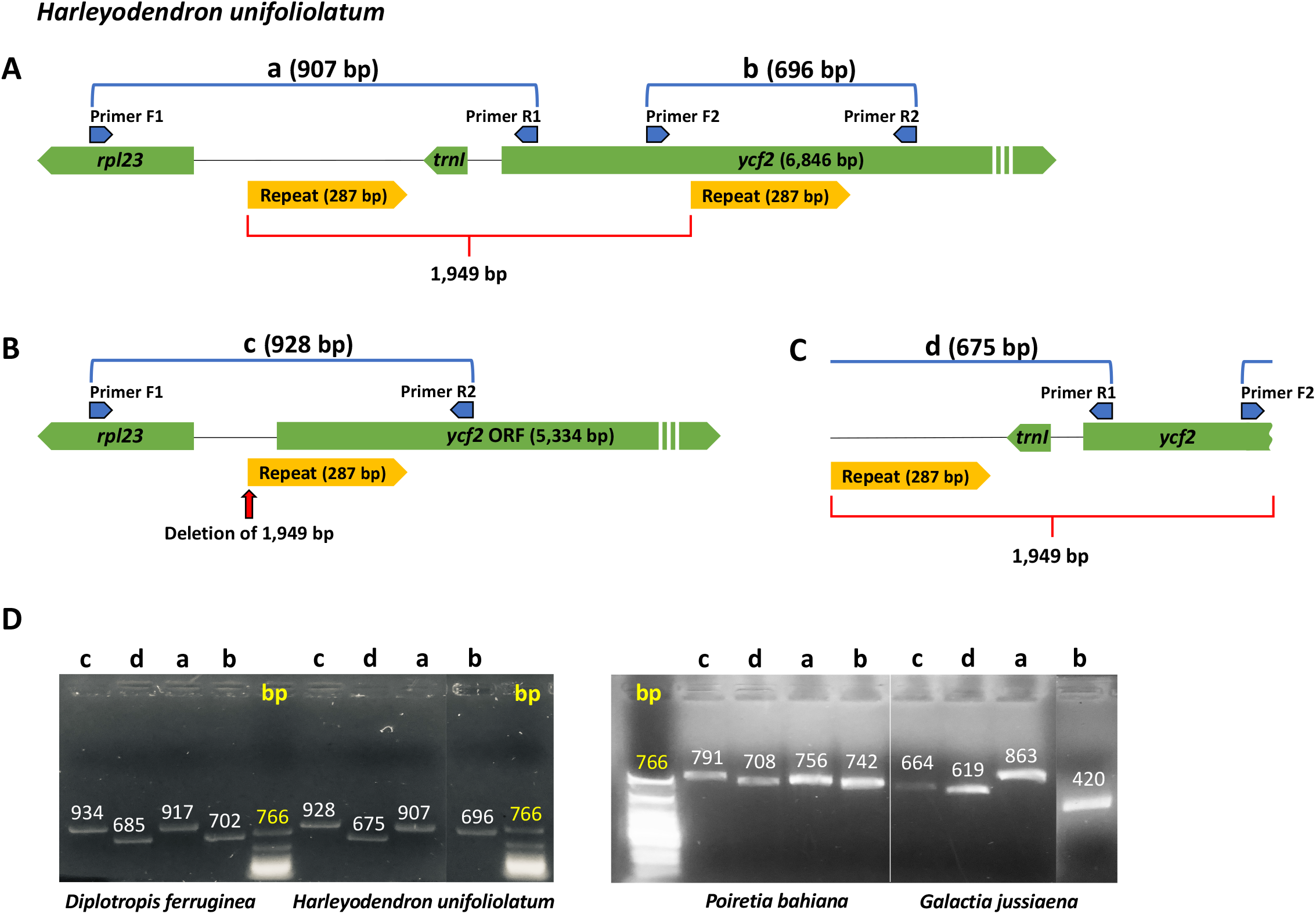
Repeat-mediated, intraindividual plastome haplotype variation. Schematic diagrams represent the region of interest in the assembled plastome of *Harleyodendron unifoliolatum* (A), the same region with a hypothesized deletion of 1949 base pairs (bp; B) and the excluded 1949 bp fragment (C). Labeled green, yellow and blue block arrows indicate size and orientation of genes, repeats and amplification primers, respectively. Primer sequences are given in Table S2. Amplification products for four taxa (D) were separated by electrophoresis along with basepair (bp) size marker (lanes labeled bp). Letters above lanes indicate the relevant predicted products in A, B and C.

Direct repeats could facilitate sequence deletion, therefore plastome assemblies for four taxa representing a range of intact repeat sizes were edited to remove the region of sequence between R1 and R2 along with one copy of the repeat (Figure 3A & 3B) and all quality-filtered paired-end reads were mapped to the revised draft. A number of reads mapped to the abbreviated draft and this regional variation was confirmed by PCR and Sanger sequencing for the four taxa *Harleyodendron unifoliolatum* (287 bp repeat), *Diplotropis ferruginea* (308 bp repeat), *Poiretia bahiana* (108 bp repeat) and *Galactia jussiaeana* (76 bp repeat)(Figure 3D). The region including the sequence between the repeats plus one repeat copy ranged from 1094 bp to 1959 bp and was extracted from the complete plastome assemblies for read mapping (Figure 3C). Several reads mapped to this fragment in *H. unifoliolatum* and *P. bahiana* and Sanger sequenced PCR amplicons were congruent with the expected regions in plastome drafts and the hypothesized fragments (Figure 3D). In all examined cases the *ycf2* ORF was maintained.

## Discussion

Since the time of Mendel and his peas (see Abbot and Fairbanks, 2016) legumes have been employed as model systems to explore questions across the varied disciplines of basic and applied biological research. Legumes, i.e. members of the rosid family Fabaceae, are widely distributed and represent the third largest of all angiosperm families. Representing ∼7% of angiosperm species, it would be difficult to overstate the importance of Fabaceae, not only as a facet of natural ecosystems, but also as a source of food, timber, medicines and forage. Furthermore their role in atmospheric nitrogen fixation makes legumes extremely valuable in both natural and agricultural settings (Graham and Vance, 2003). Virtually all dietary legumes are derived from a single subfamily, Papilionoideae, the largest of the six Fabaceae subfamilies. Also among them is the alfalfa congeneric and important model species *Medicago truncatula*.

Despite their importance, generations of study have not yet resolved the relationships among the 22 recognized clades in the subfamily (Cardoso *et al*., 2012, 2013). For phylogenetic inference, extensive sampling at deep nodes was undertaken providing an opportunity to compare plastome evolution across this fascinating and diverse subfamily. Here, 33 newly sequenced papilionoid plastomes were combined with plastomes of previously sequenced taxa to evaluate and compare changes in plastome structure. Noted were a number of inversions, gene and intron losses and a few instances of LSC tandem repeat accumulation. For the most part, gene and intron losses were previously reported for different legume taxa/lineages (Guo *et al*., 2007; Cai *et al*., 2008; Jansen *et al*., 2008; Magee *et al*., 2010; Tangphatsornruang *et al*., 2010; Sabir *et al*., 2014).

The pseudogenization and loss of *rpl20*, which is lacking in *Pterodon abruptus* (Dipterygeae), has only been reported for representatives of three subgenera of *Passiflora* (Rabah *et al*., 2018). Functional replacement by a duplicated, nuclear-encoded *rpl20* was proposed based on phylogeny, transit peptide analyses, and structural similarity to the mitochondrial-targeted copy (Shrestha *et al*., 2020). Substitution of Rpl20 in *P. abruptus* plastids could be addressed similarly, with nuclear transcriptome data. Although there were no unique intron losses uncovered, three sampled taxa have lost *clp* intron 2. Intron 1 of *clpP* was lost at the base of the IRLC (Jansen *et al*., 2008), however *Tibetia liangshanensis* was noted to have also lost *clpP* intron 2. A second species of *Glycyrrhiza, G. lepidota*, lacked *clpP* intron 2 as previously shown in *G. glabra* (Sabir *et al*., 2014). The two species are not sister (Duan et al., 2020), however expanded sampling could indicate that this is a single loss for the genus. Both *clpP* introns were missing in the IR-lacking *Camoensia scadens*.

Both unique and shared inversions were detected in sampled taxa. Among the shared inversions, was the 36 kb inversion first identified in *Lupinus luteus* (Martin *et al*., 2014) and further investigated by Schwarz et al. (2015) where the same inversion was detected in both *Lupinus alba* and the distantly related *Robinia pseudoacacia*. This LSC inversion, which lies within the larger 50 kb inversion (Doyle *et al*., 1996), was suggested to arise as a result of so-called flip-flop recombination (Martin *et al*., 2014). The inversion was likely generated by mechanisms similar to those that change the relative orientation of the single copy regions as a result of non-homologous annealing between repeats on different genome copies during replication and/or repair processes (Bendich, 2004; Maréchal and Brisson, 2010; Oldenburg and Bendich, 2015). Identification of the 36 kb inversion in *Genista* and the sister clade including *Sophora* (Sophoreae) supports the suggestion that the inversion is present in, but not restricted to, core genistoid legumes (Martin *et al*., 2014; Schwarz *et al*., 2015; Choi and Choi, 2017). Further sampling will likely reveal more instances of the 36 kb inversion in the Fabaceae as the 29 bp inverted repeat contained in *trnS* sequences was identified in plastomes of three subfamilies, Cercidoideae, Detarioideae and Papilionoideae (Schwarz *et al*., 2015). Given the potential for inversion of the sequence between the *trnS* sequences, it is plausible that its polarity shifts between alternate conformations in a given lineage.

Defined by a single genomic feature, the 50-kb inversion clade includes almost all major papilionoid lineages, except for the earliest-diverging Swartzioid, ADA, and *Styphnolobium* clades (Cardoso *et al*., 2012). This inversion occurred within the LSC in the majority of Papilionoideae and marks a major subdivision in the legume phylogeny (Doyle *et al*., 1996; Wojciechowski *et al*., 2004; Cardoso *et al*., 2012; LPWG, 2017).

Although long considered an unequivocal synapomorphy, *Sesbania grandiflora* experienced complete reversion of the involved sequence. Despite the many assembled legume plastomes (e.g. Jansen *et al*., 2008; Choi *et al*., 2019; Zhang *et al*., 2020) and wide screening for the 50 kb inversion across the family (Doyle *et al*., 1996), thus far only *S. grandiflora* has experienced reversion of the 50 kb sequence. Given the propensity of legumes to undergo major genomic rearrangements, examining additional plastomes from across the taxonomic diversity of each lineage in the family may uncover more reversion events.

Tandem and dispersed repeat accumulation varied across taxa and no obvious relationship could be drawn between dispersed repeat content and inversions, a proxy for plastome structural stability. In three species repeat accumulation exceeded 10% of plastome sequence, *Tabaroa caatingicola* (11.97%), *Trifolium repens* (11.17%) and *Hedysarum taipeicum* (18.62%). While *T. caatingicola* substantially increased LSC and plastome size (>170 kb) through accumulation of dispersed tandem repeat families, *T. repens*, like *T. subterraneum* (Milligan *et al*., 1989; Cai *et al*., 2008), accumulated more dispersed repeats as did *H. taipeicum*. Both *H. taipeicum* and *T. caatingicola* were colinear with species in sister genera, while *T. repens* was rearranged relative to *Lens culinaris* and *Wisteria*. Among the *Trifolium* species sequenced to date, all members of the ‘refractory clade’ (Sveinsson and Cronk, 2014) display moderate to massive rearrangement relative to each other and to outgroups (Cai *et al*., 2008; Sabir *et al*., 2014; Sveinsson and Cronk, 2014) suggesting that unique phenomena have influenced plastome structure in the clade as elevated content of both dispersed and tandem repeats has not influenced gene order arrangements in other legume lineages.

Most repeat accumulation, both tandem and dispersed, occurred in the LSC of IR-containing taxa. While independent IR expansions contributed to notable plastome size expansion in some taxa (Table S1), minor variances in IR size were often mediated by repeats affecting *ycf2* length. Across all IR-containing papilionoids plastome size ranged from ∼132 kb to ∼179 kb and averaged ∼158 kb. The four largest plastomes were found in *Tabaroa*, and three examples that exhibited IR expansion. Both *Ormosia* and *Hymenolobium* experienced migration of the LSC boundaries expanding IR content to include ∼41 kb and ∼36 kb of duplicated sequence, respectively. *Myroxylon* contained the largest plastome among those examined at ∼179 kb, duplicating much of the former SSC in its ∼41 kb IR.

The gene encoding ATPase motor protein Ycf2 (Kikuchi *et al*., 2018) is typically the largest plastome gene (Shinozaki *et al*., 1986; Drescher *et al*., 2000). It is, however highly variable in length and several lineages contain divergent, degraded and/or pseudogenized copies (Downie *et al*., 1994; Ruhlman and Jansen, 2018). Length variation in *ycf2* sequences can contribute dramatically to IR and overall plastome size variation given that the gene in typical plastomes ranges up to ∼8 kb (6884 bp in *Arabidopsis*; Sato *et al*., 1999). A very high frequency of deletions has been noted in the 5’ region of *ycf2*. Both typical (i.e. *Spinacia*) and atypical (i.e. *Pelargonium*) plastomes contain large deletions in *ycf2* and a survey of 279 species revealed deletions of 200 to 500 bp in representatives of 17 families (Downie *et al*., 1994). High identity direct repeats in the *ycf2* 5’ coding region were identified across papilionoid legumes beginning with the *Andira* clade representative, *Hymenolobium*. Ranging in size from 76 bp to 308 bp, the repeats were detected in all taxa that have not experienced IR loss and repeat size distribution had a strong phylogenetic signal (Figure S1, Table S1). In most IR-lacking taxa, the same repeats were found to be degraded, perhaps associated with the reduction in gene conversion templates following IR loss. Interestingly, interaction between these direct repeats has likely given rise to intraindividual length polymorphism in the gene and generated alternative plastome haplotypes. Read mapping and Sanger sequences of PCR amplicons support the presence of up to three plastome haplotypes in the four taxa shown in Figure 3 (also see Table S4).

The detection of different plastome haplotypes within individuals, other than single copy orientation isomers (Palmer, 1983), has dispelled the long-held notion that the wild-type plastome is uniformly homoplasmic (Guo *et al*., 2014; Ruhlman *et al*., 2017; Lee *et al*., 2020). Given that a proportion of the plastome exists in a complex and branched state, and has the possibility for strand annealing to occur at homeologous sites, the potential for alternate arrangements to accumulate should be fairly intuitive. While it is no great challenge to grasp the presence of two haplotypes, one containing a full length *ycf2* and one lacking the 5’ end and the upstream *trnI* gene, it is more difficult to imagine a scenario that allows for the maintenance and detection of the predicted deletion product, an ∼1950 bp, possibly circular subgenome in a wild-type, photosynthetic angiosperm plastome.

Read mapping and amplicon sequencing suggested the small haplotype was indeed present (Figure 3C, Table S2) and may exist as a circular molecule. Mitochondrial genomes have been shown to contain subgenomic circular intermediates (Park *et al*., 2014; Gualberto and Newton, 2017) and minicircles containing one to three genes of ∼2 to 3 kb in size have been detected in Peridinin-containing dinoflagellate plastomes (Zhang *et al*., 1999; Barbrook *et al*., 2006) whose replication involves rolling circle amplification (Leung and Wong, 2009). Following genetic transformation, monomeric plasmids persist briefly in *Chlamydomonas* plastids (Boynton *et al*., 1988) and an unstable DNA minicircle of 868 bp was maintained for several months following plastid transformation in *Nicotiana tabacum* (Staub and Maliga, 1994). Aberrant amplification via a rolling circle mechanism would generate a concatemer of the fragment that would map as a circle, as was suggested for entire plastomes (Kolodner and Tewari, 1975; Bendich 2004). Given the highly complex branching structures detected in several angiosperm plastomes, it is also plausible that the detected ‘minicircle’ represents a branch point or a replication intermediate. Furthermore, nuclear or mitochondrial sequence reads could be supporting the hypothesized junction, given the limited read depth relative to that of the complete unit genome arrangement (Table S1, S3).

Instances of IR loss are accumulating as more diverse taxa are sequenced, and IR-lacking taxa have been identified in several unrelated lineages (Lavin *et al*., 1990; Guisinger *et al*., 2011; Wu *et al*., 2011; Sanderson *et al*., 2015; Barrett *et al*., 2016; Zhang *et al*., 2016; Cauz-Santos *et al*., 2020; Jin *et al*., 2020a,b). Only two reports currently describe the apparent reappearance of the IR in clades thought to lack this feature, the *Erodium* long branch clade (Blazier *et al*., 2016a) and in the IRLC legume *Medicago minima* (Choi *et al*., 2019). The lack of the IR was once considered a synapomorphy for the clade that includes *M. minima*, and with the sequencing of the *Camoensia scadens*, a non IRLC papilionoid, IR loss can no longer be considered an exclusive character.

Like IRLC plastomes, the arrangement of genes flanking the remaining repeat copy in *Camoensia* suggest the loss of what is commonly referred to as IRa (Shinozaki *et al*., 1986). Recognizing that both the LSC and the SSC invert their polarity depending on annealing during replicative recombination between IR copies in different plastome monomers (Maréchal and Brisson, 2010; Oldenburg and Bendich, 2015), it may be more appropriate to define the orientation in which the single copy regions were arranged at the time of the deletion. For example, in the *Camoensia* plastome (Figure 2), the genes *ndhF* and *trnH-GUG* would be adjacent on the same molecule (head-to-tail arrangement of monomeric units; (Oldenburg and Bendich, 2004), suggesting that the two copies included in the IR were reduced to one copy while the single copy regions were in the orientation that placed these sequences at opposite IR boundaries. Here the LSC was in the orientation that is most commonly diagrammed for sequenced angiosperm plastomes (*N. tabacum trnH* begins at nucleotide 6; Shinozaki *et al*., 1986), while the SSC was in opposite orientation. Of course, if the SSC was oriented as typically depicted, i.e. *ndhF* situated at or near the junction with IRb, while the LSC was in reverse orientation, the same adjacency would result from the loss of ‘IRb’ rather than ‘IRa’. This serves as a reminder that the designation of IR copies as ‘a’ or ‘b’ is a semantical convenience observed for discussion. In addition to orientation changes, it is likely that in a plastome concatemer, there will be multiple adjacencies depending on junctions between genome units (Oldenburg and Bendich, 2004; 2016).

Most commonly, plastomes are depicted with the *rpl23* transcription unit intact, regardless of IR presence or absence. It was suggested that ‘IRa’ may be more dispensable than ‘IRb’ as loss of the latter would disrupt the *rpl23* transcription unit (Blazier *et al*., 2016a), however among a predicted population of isomers, selection should favor replicative amplification of the isoform that preserves its transcription capacity. The same could be said regarding the orientation of the SSC relative to *ycf1*. As more plastomes are elucidated there will be more losses of the typical IR noted. In addition to Pinaceae (Wu *et al*., 2011), *Drypetes* (Malpigiaceae; Jin *et al*., 2020a,b), *Passiflora* (Section *Xerogona*, Passifloraceae; Cauz-Santos *et al*., 2020) and possibly two species of *Mammilaria* (Cactaceae, Solórzano *et al*., 2019) as well as *Tahina* (Chuniophoeniceae; (Barrett *et al*., 2016) appear to have lost IRb while *Erodium, Carnegia* and IRLC plastomes were assembled in such a way as to suggest the loss of IRa. Accumulating evidence suggests that there may be no explicit functional utility with respect to which of the two sequences repeated in the typical plastome IR is retained.

IR expansions and contractions that perpetuate repeats flanking IR sequences could promote IR loss. Expansion in *Myroxylon* and *Hymenolobium* duplicated the sequence at *rpl16* across the IR, in *Camoensia* retraction of the (hypothetical) IR boundary may have left the copy in place adjacent to *ndhF* (Figure 2). Recombination between nonhomologous loci, sometimes referred to as illegitimate recombination, would have the potential to exclude the IR copy contained by the repeats. Notably, the tandem repeat accumulation in the lineages that lead up to *Camoensia* was not present in the contemporary *Camoensia* plastome, nor in any subsequent lineages until *Trifolium* (Figure 1) and intergenic regions were concomitantly smaller. Blazier *et al*. (2016b) proposed a period during which proteins that suppress illegitimate recombination are compromised allowing repeats to proliferate. Restoration of expression or the spread of alleles that better suppress illegitimate recombination halts this potentially deleterious process.

Dispersed repeats may have played a role in IR deletion suggesting that IR loss resulted, directly or indirectly, from other factors causing overall genome lability. With a range of IR phenomena for comparative genomics, broader sampling across the papilionoid system may help to unravel a persistent question: What contribution do the IR and non-IR repeats play in plastome structural dynamics? This remains a ‘chicken or the egg’ question with respect to plastome evolution.

## Acknowledgements

This work was supported by grants from the National Science Foundation (DEB-1853024) to R.K.J., T.A.R. and M.F.W., the Texas Ecological Laboratory (EcoLab) to R.K.J., T.A.R. and I.C., the Sidney F. and Doris Blake Professorship in Systematic Botany to R.K.J., the CNPq (Research Productivity Fellowship no. 308244/2018-4; Universal no. 422325/2018-0) and FAPESB (Universal no. APP0037/2016) to DC. The authors thank the TEX-LL, HUEFS, and RB herbaria for voucher deposition and the Desert Legume Program for seeds; Alessandra Schnadelbach and Hedina Basile for their support at UFBA. The authors thank Luciano Paganucci de Queiroz (UEFS) and George Yatskievych (TEX/LL) for arranging a formal Material Transfer Agreement (Decree number 8,772) to facilitate research activities between the institutions.

---

**Figure S1: Maximum likelihood phylogeny of Papilionoideae**. The phylogeny is based on 76 shared plastome genes (Table S3) and includes 64 legume taxa (59 Papilionoideae) and the outgroups *Polygala fallax* (MT762166), *Suriana maritima* (NC_047313) and *Quillaja saponaria* (NC_047356). Clades are highlighted with shaded color and papilionoid clades are named according to Cardoso *et al*. (2012). Bootstrap values less than 100% are shown at the nodes. The size of conserved repeats around *ycf2* is given in basepairs (bp, right) for each papilionoid taxon (see Figure 3) that contained them. The most common repeat size was 287 bp and its distribution is indicated with values in black font (see also Table S1. The scale bar indicates number of substitutions per site and branch tips are extended by broken lines to taxon name. IRLC, inverted repeat-lacking clade.

**Figure S2: Whole genome alignment of *Camoensia scadens***. The inverted repeat (IR) lacking *Wisteria floribunda* was used as a reference. Syntenic regions are indicated by colored, locally collinear blocks (LCBs). Histograms inside each LCB represent pairwise nucleotide sequence identity. Inversions are shown as blocks flipped across the plane. Ribosomal RNA genes are colored red.

**Table S1: Plastome sequencing statistics**.

**Table S2: Amplification and sequencing primers employed to investigate alternative plastome haplotypes**.

**Table S3: Shared plastome protein coding genes included in phylogenetic analysis**.

**Table S4: Reads mapped to alternative haplotypes**.

