## Supplementary figures and images for "The chicken or the egg? Plastome evolution and a novel loss of the inverted repeat in papilionoid legumes"

### Supplemental Figure S1

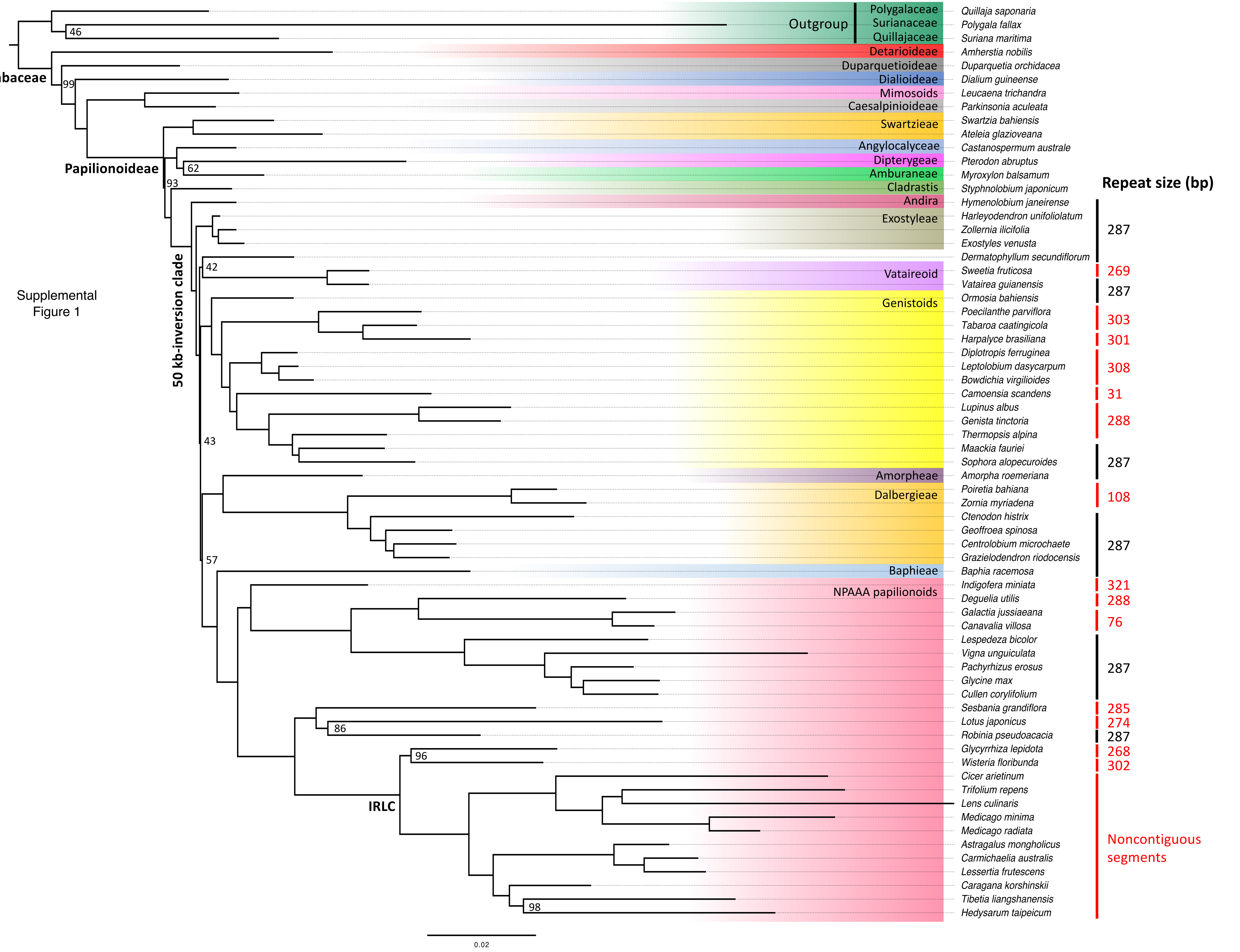


Supplemental Figure 1

### Supplemental Figure S2

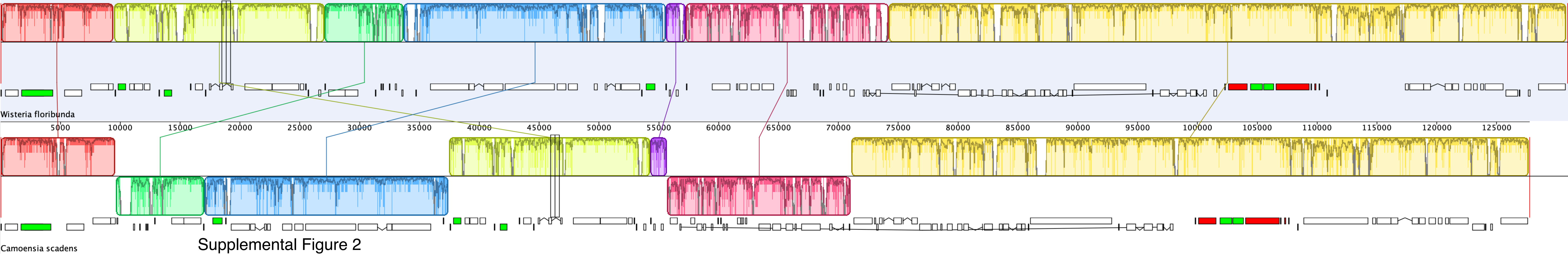


Supplemental Figure 2
