## Supplemental Table S2 for "The chicken or the egg? Plastome evolution and a novel loss of the inverted repeat in papilionoid legumes"

**Table S2:** Amplification and sequencing primers employed to investigate alternative plastome haplotypes.

| **Primer name** | **Primer sequence (5'-3')** | **Primer length (nt)** |
| --- | --- | --- |
| MG04-1-F | AGGTCGCATTCTTCTACCCT | 20 |
| MG04-1-R | ATCATTCCCGCAAGAGGTCC | 20 |
| MG04-2-F | TGCCGACTATTCATGGAACG | 20 |
| MG04-2-R | CCGCTGAATTGAATTGGGTCC | 21 |
| BA08-2-F | GGAGCGATTCCCTAAATGCC | 20 |
| BA13-1-F | TCGGTCGCACTTTTCTACCC | 20 |
| BA13-1-R | TGGTGTAACGTGTATCCCCC | 20 |
| BA13-2-F | CATGGAACGTGAGAAGGAGATG | 22 |
| BA13-2-R | CCGCGGAATTGAATTGGGTC | 20 |
