## Supplemental Table S3 for "The chicken or the egg? Plastome evolution and a novel loss of the inverted repeat in papilionoid legumes"

**Table S3:** List of 73 plastid protein-coding genes used for the phylogenetic analysis.

|  | **Gene** |
| --- | --- |
| **Photosynthesis** | *atpA, atpB, atpE, atpF, atpH, atpI, ndhA, ndhB, ndhC, ndhD, ndhE, ndhF, ndhG, ndhH, ndhI, ndhJ, ndhK, petA, petB, petD, petG, petL, petN, psaA, psaB, psaC, psaI, psaJ, psbA, psbB, psbC, psbD, psbE, psbF, psbH, psbI, psbJ, psbK, psbL, psbM, psbN, psbT, psbZ, rbcL* |
| **Genetic system** | *matK, rpl2, rpl14, rpl16, rpl23, rpl32, rpl36, rpoA, rpoB, rpoC1, rpoC2, rps2, rps3, rps4, rps7, rps8, rps11, rps12, rps14, rps15, rps18, rps19* |
| **Other** | *ccsA, cemA, clpP, ycf1, ycf2, ycf3, ycf4* |
