## Supplemental Table S4 for "The chicken or the egg? Plastome evolution and a novel loss of the inverted repeat in papilionoid legumes"

**Table S4:** Reads supporting alternative haplotypes

| **Species** | **Repeat size** | **Mapped read pairs** | |
| --- | --- | --- | --- |
|  |  | **Alternative type** | **Fragment of ~2 kb** |
| *Harleyodendron unifoliolatum* | 287 | 2 | 20 |
| *Diplotropis ferruginea* | 308 | 7 | 39 |
| *Poiretia bahiana* | 108 | 2 | 5 |
| *Galactia jussiaeana* | 76 | 5 | 16 |
